# Stable Engraftment of a Human Gut Bacterial Microbiome in Double Humanized BLT-mice

**DOI:** 10.1101/749093

**Authors:** Lance Daharsh, Amanda E. Ramer-Tait, Qingsheng Li

## Abstract

**Background:** Humanized mice featuring a functional human immune system are an important pre-clinical model for examining immune responses to human-specific pathogens. This model has been widely utilized to study human diseases that are otherwise impossible or difficult to investigate in humans or with other animal models. However, one limitation of using humanized mice is their native murine gut microbiome, which significantly differs from the one found in humans. These differences may be even greater for mice housed and bred in specific pathogen free conditions. Given the importance of the gut microbiome to human health and disease, these differences may profoundly impact the ability to translate the results from humanized mice studies to human disease. Further, there is a critical need for improved pre-clinical models to study the complex *in vivo* relationships of the gut microbiome, immune system, and human disease. We therefore created double humanized mice with both a functional human immune system and stable human-like gut microbiome.

**Results:** Surgery was performed on NOD.*Cg-Prkdc^scid^II2rg^tm1Wjl^*/SzJ (NSG) mice to create bone-marrow, liver, thymus (BLT) humanized mice. After immune reconstitution, mice were treated with broad spectrum antibiotics to deplete murine gut bacteria and then transplanted with fecal material from healthy human donors. Characterization of 173 fecal samples obtained from 45 humanized mice revealed that double humanized mice had unique 16S rRNA gene profiles consistent with those of the individual human donor samples. Importantly, transplanted human-like gut microbiomes were stable in mice for the duration of the study, up to 14.5 weeks post-transplant. Microbiomes of double humanized mice also harbored predicted functional capacities that more closely resembled those of the human donors compared to humanized mice.

**Conclusions:** Here, we describe successful engraftment of a stable human microbiome in BLT humanized mice to further improve this preclinical humanized mouse model. These double humanized mice represent a unique and tractable new model to study the complex relationships between the human gut microbiome, human immune system, and human disease *in vivo*.

## Background

The complex ecosystem of the gut microbiome plays a critical role in human health and disease [1–5]. Specifically, the gut microbiome has a highly reciprocal and dynamic relationship with the immune system. Antigens derived from the gut microbiome influence host immune responses, and the immune system in turn contributes to shaping the spatial distribution and composition of the gut microbiota [6–8]. Humanized mice (hu-mice) with an engrafted human immune system have facilitated important advancements in the study of human cancer, autoimmune diseases, hematopoiesis, and infectious diseases [9–17]. However, gut microbiomes of hu-mice are murine in origin and are often not well-characterized in translational studies. The murine gut microbiome differs substantially in composition and function from that of humans [18], primarily due to anatomical differences as well as other factors such as diet [19]. Considering the importance of the gut microbiota to proper immunological development and influencing immune responses, the murine origin of the microbiome harbored by hu-mice could affect translational study outcomes. Consequently, a need exists to not only characterize the gut microbiomes of hu-mice, but also impart these mice with a more human-like gut microbiome to improve the translational aspects of the model for human medicine.

The creation of a new pre-clinical hu-mice model to study the human immune in the context of a human microbiome offers numerous benefits over existing options. Many aspects of human disease are difficult or impossible to study directly in humans due to practical or ethical concerns. Non-human primate models are informative but are genetically outbred, and large studies are often resource and cost prohibitive. Many important discoveries in the microbiome field have been made using mouse models; however, translating results from mouse studies to humans has often proved difficult. The use of germ-free mice reconstituted with human-like gut microbiomes has been the gold standard in studying the relationship of the gut microbiome to human health and disease [20, 21]. However, working with or deriving germ­ free animals requires expertise and facilities that are not always available. Further, many immunodeficient mouse strains commonly used to reconstitute a human immune system, such as NOD.*Cg-Prkdc^scid^II2rg^tm1Wjl^*/SzJ (NSG), are currently not commercially available as germ-free. We therefore created a double hu-mice model featuring both a functional human immune system and a stable human-like gut microbiome under specific pathogen free (SPF) conditions. Here, we show that double hu-mice had unique 16S rRNA gene profiles based on the individual human donor sample with which they were colonized. Importantly, the transplanted human­ like microbiome was stable in the mice for the duration of the study, up to 14.5 weeks post­ transplant. Double hu-mice also harbored gut microbiomes with a more human-like predicted functional capacity compared to their hu-mice counterparts.

## Results

### Gut microbiomes of double hu-mice are distinct and more human-like compared to hu-mice

To create double hu-mice, surgery was performed on NSG mice to create bone-marrow, liver, thymus (BLT) hu-mice. Hu-mice were then pre-treated with a cocktail of broad-spectrum antibiotics and administered fecal transplants using fecal material from healthy human donors (see Methods and Daharsh et al. [22] for detailed descriptions and demonstrations). Multiple cohorts of double hu-mice were created using fecal material from one of three unique healthy human donors or an equal mixture of all three (Table 1). We used 16S rRNA gene sequencing and characterized the gut bacterial microbiome of 100 fecal samples from 16 double hu-mice and compared them with 67 fecal samples representing the pre-existing murine gut bacterial microbiomes of hu-mice and the profiles of the 4 human fecal donor samples. To visualize beta­ diversity relationships between hu-mice, double hu-mice, and the human donor samples, non-metric multidimensional scaling (NMDS) and principal coordinate analysis (PCoA) plots were created (Fig. 1). Using NMDS, the gut microbiome profiles of the three groups separated into distinct clusters (Fig. 1a). Using PCoA, the human donor samples fell within the double hu-mice hull, which was distinct from the hu-mice profiles (Fig. 1b). Both dimensionality reduction methods showed that after engraftment of a human gut microbiome, double hu-mice represented a distinct population that clustered closer to the human donor samples compared to hu-mice harboring murine gut microbiomes. Importantly, there was no reversion to hu-mice profiles post-transplant. Displaying the relatedness of the samples through a hierarchical dendrogram based on Bray-Curtis distances further confirmed the similarity of double hu-mice microbiomes to the human donor samples and differentiated them from those of hu-mice (Fig. 2).

**Table 1.** Experimental summary of double hu-mice cohorts.

| Cohort | Antibiotic start | Antibiotic duration | Number of fecal transplants | Human fecal donor | Maximum weeks post-transplant | Double hu-mice | Double hu-mice fecal samples | Hu-mice | Hu-mice fecal samples |
| --- | --- | --- | --- | --- | --- | --- | --- | --- | --- |
| Antibiotic Pilot | After immune reconstitution | 9 days | 0 | NA | 0 | 0 | 0 | 6 | 9 |
| Donor 65 Cohort 1 | After immune reconstitution | 14 days | 2 (24 & 48 hr post antibiotics) | 65 | 13.5 | 3 | 35# | 3 | 23 |
| Donor 65 Cohort 2 | After immune reconstitution | 14 days | 2 (24 & 48 hr post antibiotics) | 65 | 14.5 | 2 | 16# | 8 | 8 |
| Donor 74 | After immune reconstitution | 7 days | 2 (24 & 48 hr post antibiotics) | 74 | 6 | 3 | 17# | 3* | 3* |
| Donor 82 | After immune reconstitution | 7 days | 2 (24 & 48 hr post antibiotics) | 82 | 6 | 4 | 27# | 3* | 3* |
| Donor Mix | 3 days after hu-mice surgery | 14 days | 2 (24 & 48 hr post antibiotics) | Equal mixture of 65, 72, & 84 | 9 | 4 | 21# | 9 | 10 |
| # Includes 1 pre-fecal transplant sample for each of the double hu-mice |  |  |  |  |  |  |  |  |  |
| * Control hu-mice were the same for Donor 74 and Donor 82 groups |  |  |  |  |  |  |  |  |  |

**Figure 1.**
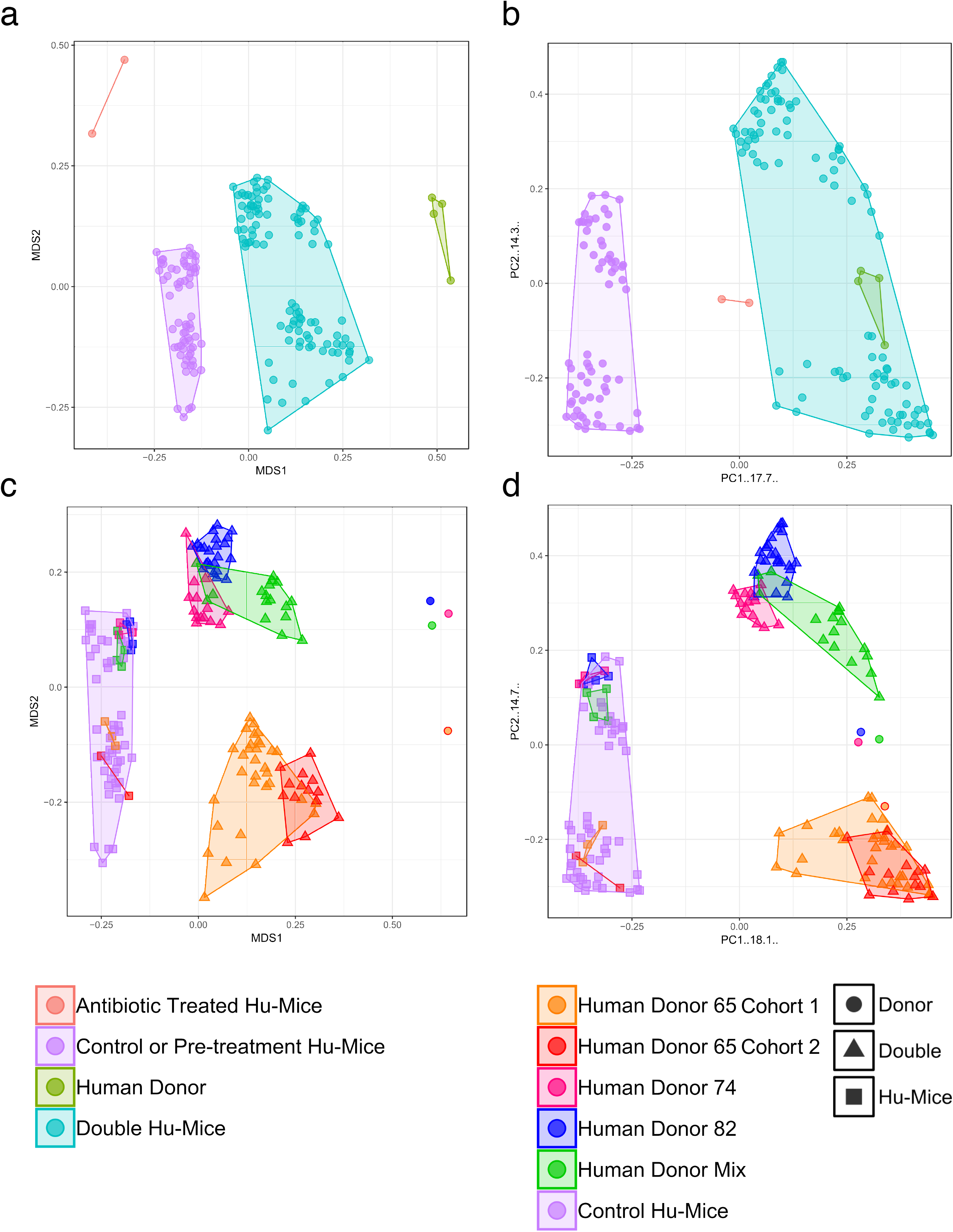
Gut microbiomes of double hu-mice are distinct and more human-like compared to hu-mice and feature donor specific profiles. A) Non-metric multidimensional scaling (NMDS) plot displaying double hu-mice as a distinct cluster between the human donor samples and pre­ treatment or untreated control hu-mice. B) Principal coordinate analysis (PCoA) plot showing the double hu-mice cluster with the human donor samples distinct from the pre-treatment or untreated control hu-mice. C) NMDS plot displaying human donor specific profiles in the double hu-mice. D) PCoA plot showing human donor specific profiles in the double hu-mice.

**Figure 2.**
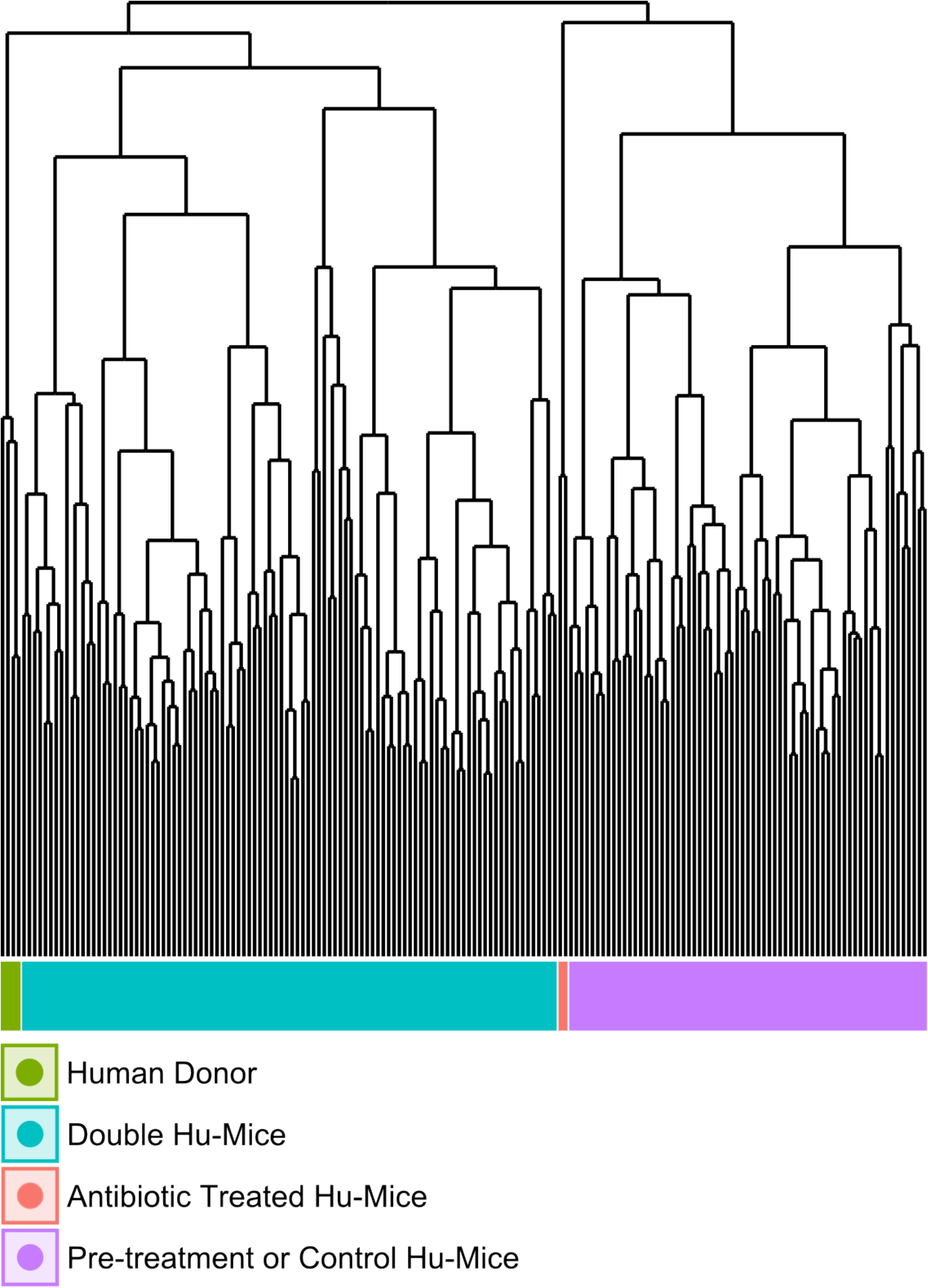
The gut microbiomes of double hu-mice cluster with that of human donor fecal samples. Dendrogram based on Bray-Curtis distances for the gut microbiome profiles of pre­ treatment and untreated control hu-mice (Pre-treatment or Control Hu-Mice), double hu-mice (Double Hu-Mice), antibiotic treated hu-mice (Antibiotic Treated Hu-mice), and human donor fecal samples (Human Donor).

We also observed that double hu-mice maintained the pre-existing relationships between gut microbiome profiles of the human fecal donors (Fig. 1c & 1d). Importantly, pre­ treatment samples from corresponding double hu-mice in each cohort had similar gut microbiome profiles as untreated control hu-mice. After engraftment, double hu-mice resembled the individual human donor that was transplanted as demonstrated by the relationships between the human donor microbiome profiles. Human donors 72 and 84 had more similar profiles to one another than to human donor 65. This relationship was maintained in the double hu-mice after engraftment. We also prepared an “un-biased” human sample by mixing equal parts of the three human donor fecal samples, designated hereafter as donor mix. The microbiome profile of this mixed sample resembled a mixture of the three individual human donor profiles. Specifically, the mixed donor sample more closely resembled individual donors 72 and 84, and this observation was mirrored in the double hu-mice engrafted with the donor mix sample.

We also investigated the impact of antibiotic treatment duration on the engraftment of human gut microbiome. The double hu-mice engrafted with human donors 74 and 82 were generated after only 7 days of antibiotic pre-treatment. Their microbiome profiles were less similar to the human donor profiles than those from cohorts pre-treated with antibiotics for 14 days prior to fecal transplant (Fig. 1c & 1d). We found the 2 weeks of antibiotic treatment was optimal for the creation of double hu-mice. Together, these results demonstrate that our approach to generating double hu-mice is reproducible and able to create hu-mice with unique 16S rRNA gene profiles based on the individual human fecal donor.

### Gut microbiomes of double hu-mice have increased levels of alpha diversity compared to hu-mice

Due to the highly reciprocal nature of the gut microbiome and immune system, we hypothesized that highly immunodeficient mice, such as NSG, would have low pre-existing gut microbiome diversity, especially when the mice were housed under SPF conditions with limited exposure to outside sources of microbes. We tested this hypothesis and found that the gut microbial diversity of double hu-mice with a functional immune system significantly increased to the levels observed in our human donor samples compared to hu-mice. Several alpha diversity measurements confirmed that hu-mice had very low measures of alpha diversity compared to our human donor samples (Fig. 3). However, after engraftment, double hu-mice had increased species richness compared to hu-mice (P<.001) and did not differ significantly from the human donor samples (Fig. 3a). Further, the Shannon index values of the human donor samples were significantly higher than hu-mice (P<.05) but were not significantly different from the double hu-mice samples. Double hu-mice had increased Simpson index values compared to hu-mice, but the human donor samples were significantly higher than both the double hu-mice (P<.05) and hu-mice (P<.05). As expected, samples taken during antibiotic treatment had the lowest measures of all three diversity metrics tested. Alpha diversity metrics were also measured based on the human donor sample engrafted (Supplemental Figure - Alpha_Diversity.pdf). Double hu-mice engrafted after 14 days of antibiotics (donor 65 cohorts 1 & 2, donor mix) had higher levels of alpha diversity compared to double hu-mice engrafted after only 7 days of antibiotics (donors 74 & 82). Overall, double hu-mice had increased alpha diversity compared to hu-mice that was more similar to the levels observed in the human donor samples. The shorter antibiotic treatment duration (7 days versus 14 days) was associated with lower alpha diversity measurements in double hu-mice engrafted with human donors 74 or 82.

**Figure 3.**
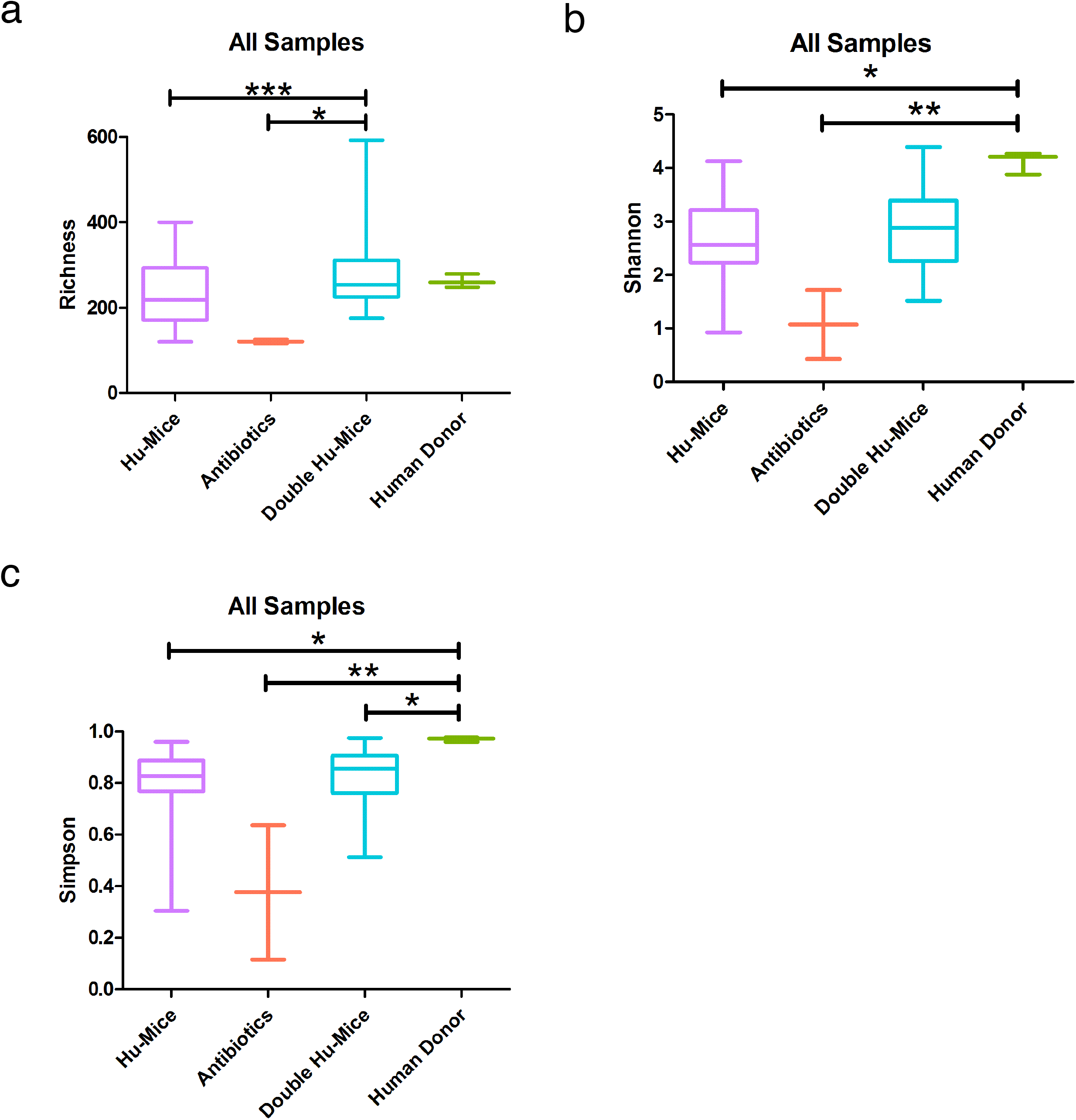
The gut microbiomes of double hu-mice have increased alpha diversity measures compared to that of pre-treatment or untreated control hu-mice. A) Species richness, B) Shannon index, and C) Simpson index. Data are shown for pre-treatment or untreated control hu-mice (Hu-Mice), antibiotic treated mice (Antibiotics), double hu-mice (Double Hu-Mice), and human donor fecal samples (Human Donor).

### The relative abundance of gut bacteria in double hu-mice is similar to that found in human donor samples

We next hypothesized that if bacteria from the human donors were successfully engrafted into double hu-mice, then we would see more taxonomic similarities between human donor gut microbiome profiles and double hu-mice versus hu-mice. Several differences were observed in the relative abundances of double hu-mice based on the length of antibiotic treatment, the human fecal donor sample engrafted, and mouse cohort (Supplemental Figure-Relative_Abundances.pdf). At the Phylum level, both double hu-mice and hu-mice samples were largely represented by *Actinobacteria, Bacteroidetes, Firmicutes,* and *Verrucomicrobia.* Interestingly, we found that hu-mice had high proportions of *Verrucomicrobia* that remained high even after antibiotic treatment and engraftment with human donor samples that had low abundances of *Verrucomicrobia.* At the Family level, hu-mice samples had a higher relative abundance of S24-7than double hu-mice and human donor samples (Fig. 4). Both hu-mice and double hu-mice samples had higher relative abundances of *Lactobacillaceae* and *Verrucomicrobiaceae* than human donor samples. The human donor and double hu-mice samples had higher relative abundances of *Bacteroidaceae* than hu-mice samples. The human donor samples also had a much higher relative abundance of *Lachnospiraceae*.

**Figure 4.**
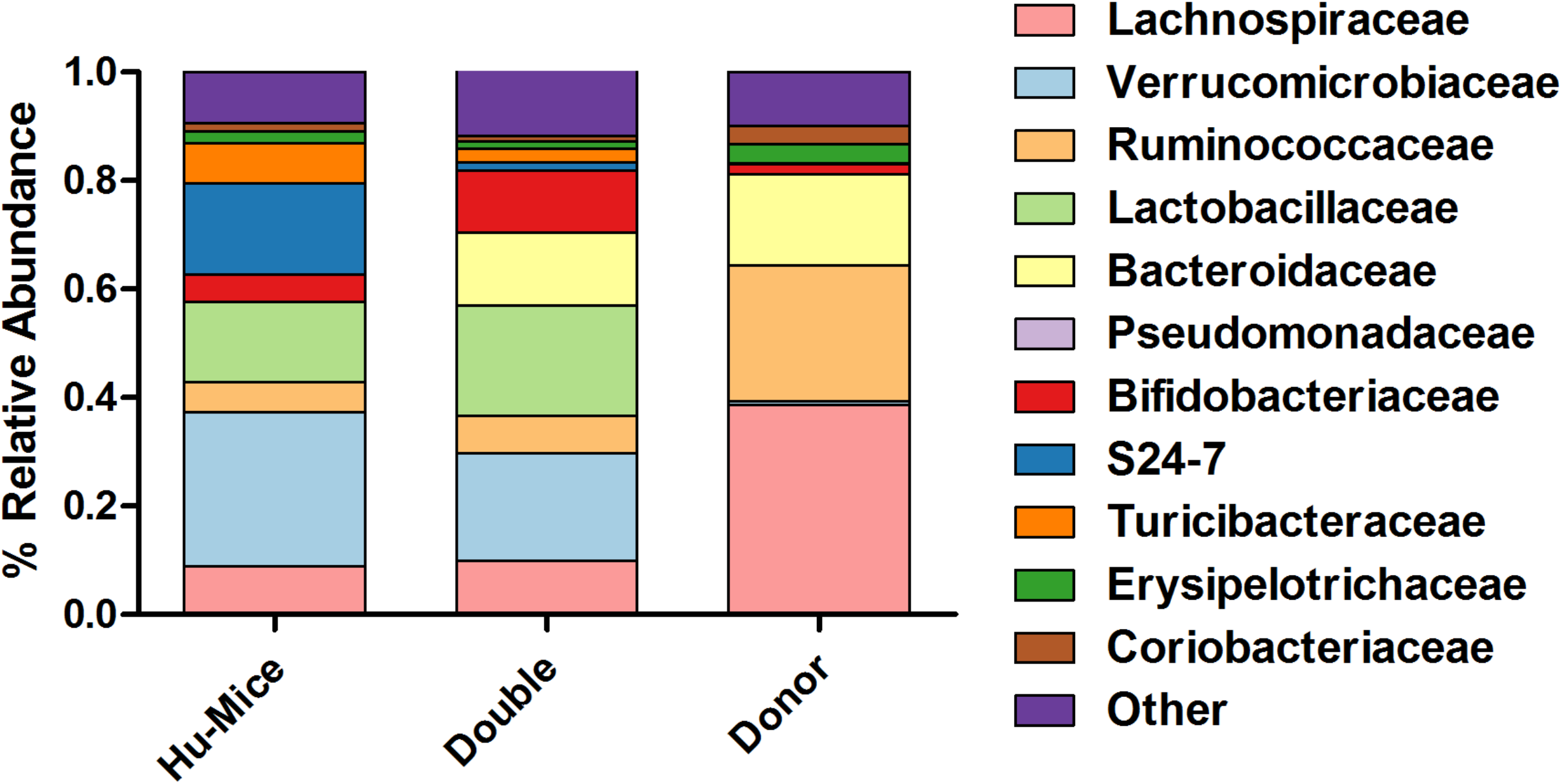
Comparison of relative abundance of taxa grouped by Family. The 11 most abundant taxa by relative percent abundance grouped by Family are shown for pre-treatment and untreated control hu-mice (Hu-mice), double hu-mice (Double), and human fecal donor samples (Donor).

We next compared the taxonomic differences between the microbiomes using Kruskal­ Wallis testing with FDR adjusted P values below .05 (Supplemental File - KW_Testing.xlsx). We found 195 significant differences between the hu-mice and human donor samples and 170 significant differences between hu-mice and double hu-mice. However, only 108 significant differences were observed between double hu-mice and human donor samples. Both the human donor and double hu-mice samples had significantly higher relative abundances of the genus *Bacteroides,* while hu-mice had a significantly higher relative abundance of the family *524-7.* Human donor samples had a higher relative abundance of the Phylum *Firmicutes* compared to both hu-mice and double hu-mice. Human donor samples had lower relative abundances of the class *Bacilli* and family *Lactobacillaceae* compared to both hu-mice and double hu-mice. Human donor samples also had a higher relative abundance of the class *Clostridia* than both hu-mice and double hu-mice samples. Significant differences found within this class were the family *Lachnospiraceae,* genus *Blautia,* genus *Roseburia,* family *Ruminococcaceae,* genus *Ruminococcus,* and species *Faecalibacterium prausnitzii.* Some of these taxa do increase in abundance in double hu-mice as compared to hu-mice, as observed with significant differences in the genus *Blautia* and genus *Ruminococcaceae.* Levels of the species *Akkermansia muciniphila* are significantly higher in hu-mice and double hu-mice samples compared to human donor samples. However, double hu-mice have significantly less relative abundance of this species than hu-mice samples.

To further determine differences in relative abundances among microbiomes from the various treatments, we used Linear discriminant analysis Effect Size (LEfSe) with a P value < .OS and LDA score> 2 (Supplemental File - LefSe.xlsx)[23]. Taxa with LDA scores higher than 4 were plotted to show significant differences between double hu-mice and human donor samples and double hu-mice and hu-mice samples (Fig. 5). Human donor samples were associated with higher relative abundances of several types of *Clostridia* including *Lachnospiraceae, Blautia, Coprococcus, Roseburia, Facalibacterium,* and *Ruminococcus* compared to double hu-mice, while double hu-mice samples were associated with *Lactobacillus* and *Akkermansia muciniphila* (Fig. 5a). Double hu-mice were associated with higher relative abundances of *Bacteroides* and several types of *Clostridia* including *Blautia, Coprococcus,* and *Ruminococcaceae,* while hu-mice samples were associated with *524-7* and *Mogibacteriaceae* (Fig. 5b). By characterizing the relative bacterial abundance in double hu-mice, we demonstrated that certain taxa, like members of *Bacteroides,* readily engrafted while others such as *Clostridia* were more difficult to transplant. Further, several species such as those found in the phylum *Verrucomicrobia* were highly prevalent in hu-mice, and antibiotic treatment followed by fecal transplant did not fully diminish or replace this population based on relative abundances. Altogether, these results demonstrate that engraftment of human donor samples significantly changed the taxonomic profile of double hu-mice and that their human-like gut microbiomes were statistically more similar to human donor profiles compared to hu-mice.

**Figure 5.**
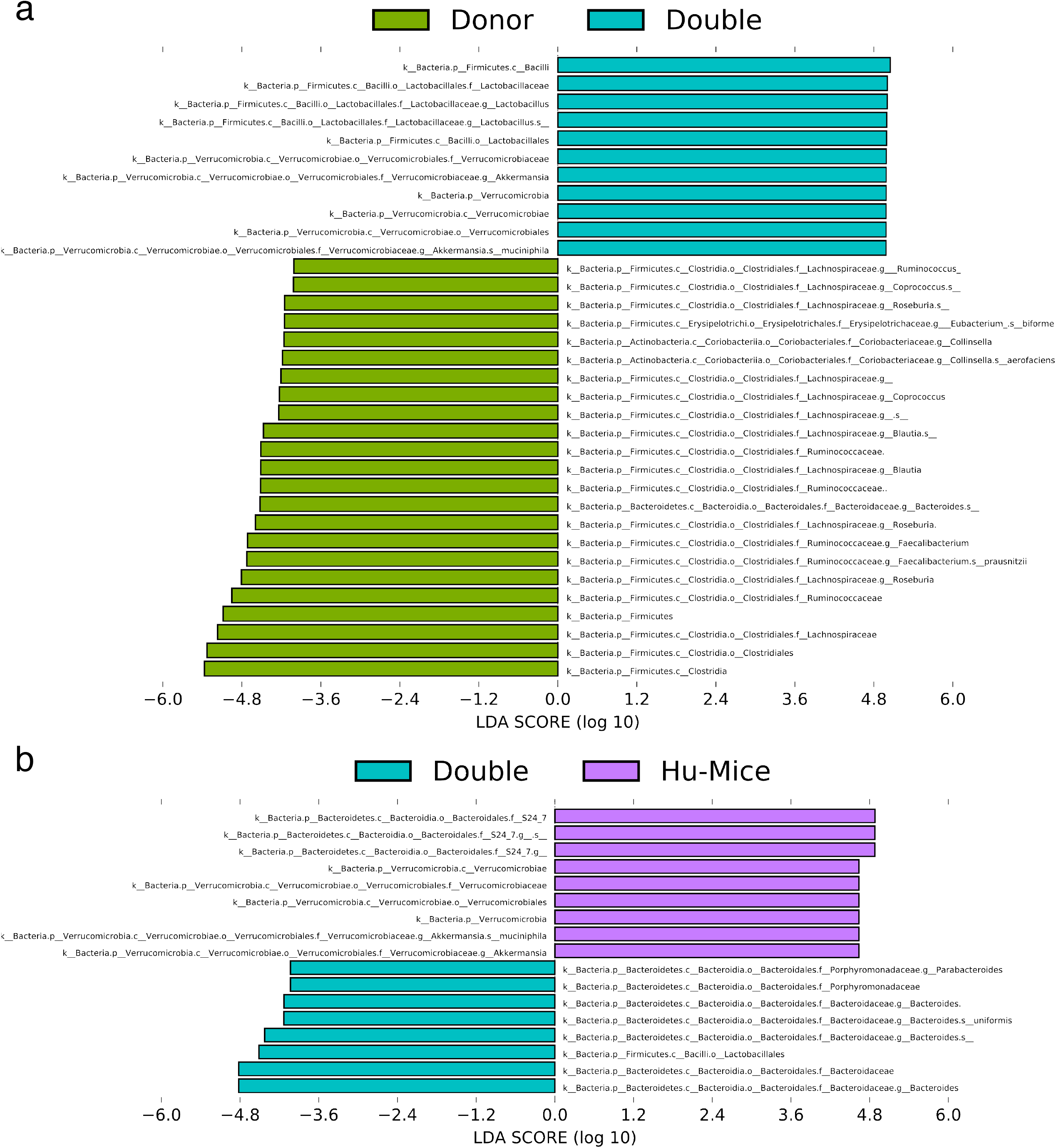
Engraftment of human fecal donor bacteria in double hu-mice as shown by LEfSe. A) All significant features with a linear discriminant analysis (LDA) score> 4.0 between human fecal donor samples (Donor) and double hu-mice (Double). B) All significant features with an LDA score> 4.0 between double hu-mice (Double) and pre-treatment or untreated control hu­ mice (Hu-Mice).

### The engrafted human-like gut microbiome in double hu-mice is stable

To evaluate the stability of the engrafted human-like gut microbiome in double hu-mice after fecal transplant, the proportion of shared amplicon sequence variants (ASVs) with the human donor sample was calculated (Fig. 6). After engraftment, double hu-mice had increased proportions of shared ASVs with their respective human donor samples, and those proportions remained higher than pre-treatment and control levels for the duration of the study. The first cohort of double hu-mice engrafted with human donor 65 had an average shared ASV proportion of 13.70% after transplant, while the pre-treatment samples had 1.06% and control samples levels had 1.42% (Fig. 6a). This increased proportion of shared ASVs was maintained for the duration of the study, up to 14 weeks post-transplant. A second cohort of double hu-mice engrafted with human donor 65 was created and a similar increase in the proportion of shared ASVs was observed (Fig. 6b). The average proportion of shared ASVs was 17.65% post­ transplant, while the pre-treatment samples had 2.60% and the control samples had 0.96%. This increased proportion of shared ASVs was maintained for the duration of the study, up to 14.5 weeks post-transplant. Double hu-mice transplanted with the mixture of all three human donors had an average shared ASV proportion of 10.91% after transplant, while the pre­ treatment samples had 0.31% and the control samples had 0.61% (Fig. 6d).

**Figure 6.**
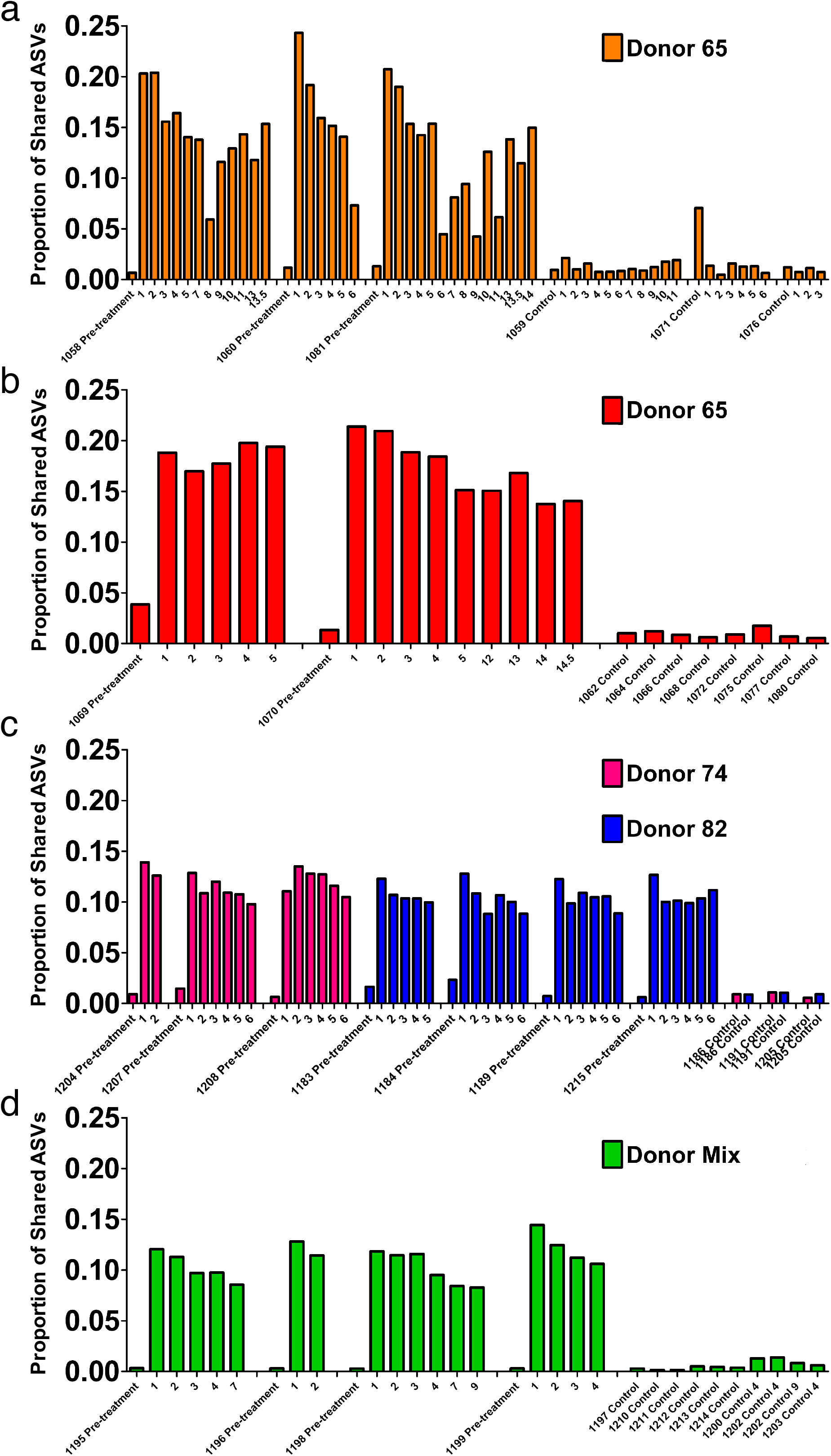
Engraftment and stability of a human-like gut microbiome in double hu-mice as determined by shared amplicon sequence variants (ASVs). A) Proportion of shared ASVs with the donor in the first cohort of double hu-mice created using fecal material from human donor 65 (Donor 65). B) Proportion of shared ASVs with donor in the second cohort of double hu-mice created using fecal material from human donor 65 (Donor 65). *C)* Proportion of shared ASVs with donor in in double hu-mice created using fecal material from human donor 74 (Donor 74) or 82 (Donor 82). D) Proportion of shared ASVs with donor in double hu-mice created using a mixture of fecal material from all three human donors (Donor Mix). X-axis numbers represent number of weeks post fecal transplant.

The proportion of shared ASVs was then calculated for cohorts of double hu-mice only treated with 7 days of antibiotics prior to fecal transplant. Double hu-mice engrafted with human donor 74 had an average shared ASV proportion of 11.85% after transplant, while the pre-treatment samples had 1.02% and the control samples had 0.86% (Fig. 6c). Double hu-mice transplanted with human donor 82 had an average shared ASV proportion of 10.56% after transplant, while the pre-treatment samples had 1.33% and the control samples had 0.95% (Fig. 6c).

To further evaluate the stability of the engrafted human-like microbiome after transplant into double hu-mice, the contributions of the human donor and pre-treatment sample to the post-transplant samples were determined using SourceTracker (Fig. 7)[24]. The first cohort of mice transplanted with human donor 65 had an average donor contribution percentage of 18.76% after transplant, while the pre-treatment samples had 0.00% and the control samples had 0.05% (Fig. 7a). At the final time point collected at 14 WPT, the donor contribution was consistent at 18.10%. The second cohort of mice transplanted with human donor 65 had an average donor contribution percentage of 29.01% after transplant, while the pre-treatment and controls samples had no donor contribution (Fig. 7b). The donor contribution percentage was 34.10% at 14.5 WPT, thus demonstrating the stable engraftment of donor bacteria. Double hu-mice engrafted with donor 74 had an average donor contribution percentage of 12.71% after transplant, while the pre-treatment samples had 0.00% and the control samples had 0.00% (Fig. 7c). At the final time point collected at 6 WPT, the average donor contribution was 18.75%.

**Figure 7.**
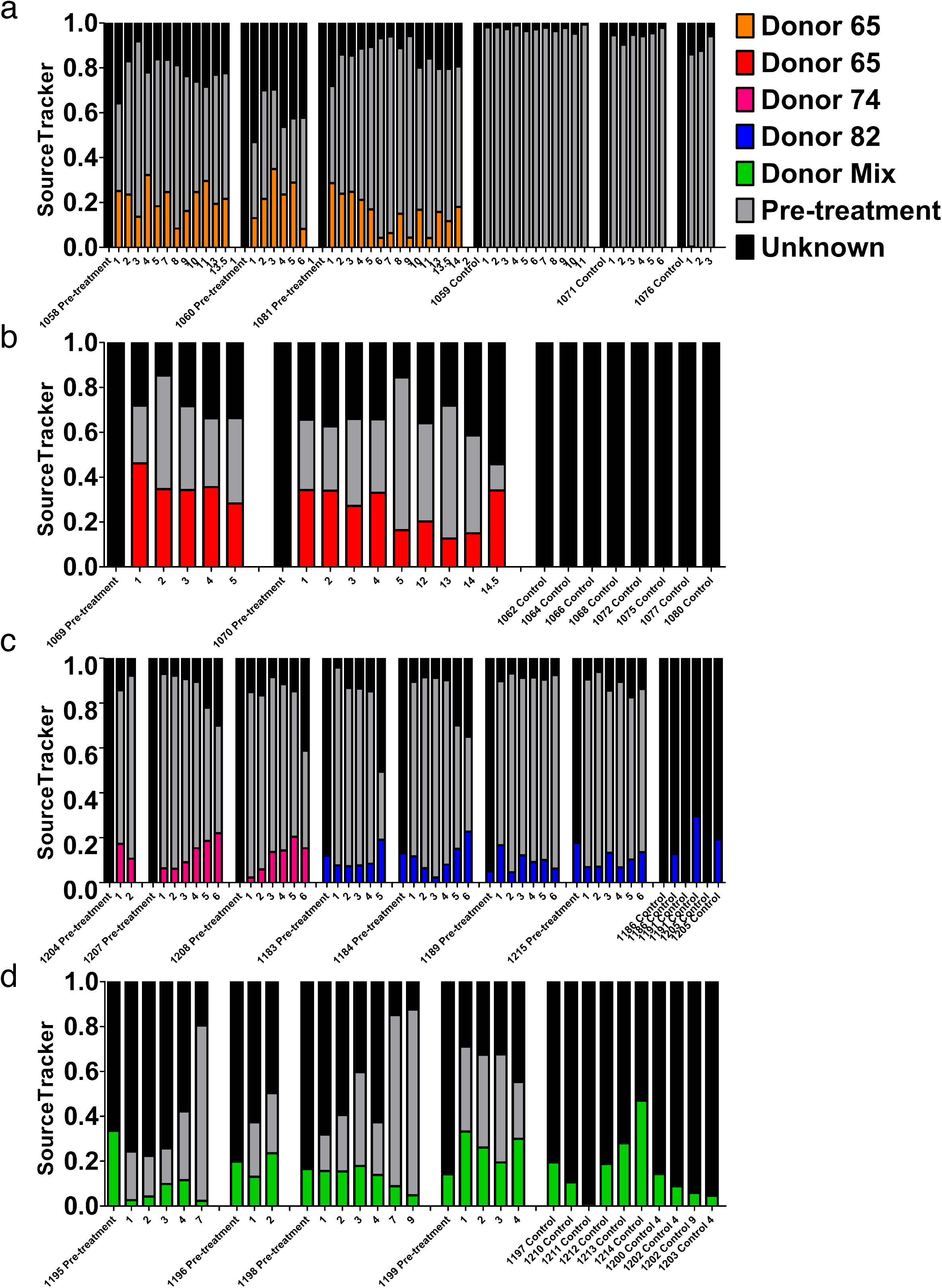
Stability of the engrafted human-like gut microbiome in double hu-mice as determined by SourceTracker. A) Contributions of the human fecal donor sample and pre­ treatment sample in the first cohort of double hu-mice created using fecal material from human donor 65 (Donor 65). B) Contributions of the human fecal donor sample and pre­ treatment sample in the second cohort of double hu-mice created using fecal material from human donor 65 (Donor 65). C) Contributions of the human fecal donor sample and pre­ treatment sample in double hu-mice created using fecal material from human donor 74 (Donor 74) or 82 (Donor 82). D) Contributions of the human fecal donor sample and pre-treatment sample in double hu-mice created using a mixture of fecal material from all three human donor (Donor Mix). X-axis numbers represent number of weeks post fecal transplant.

The SourceTracker algorithm was unable to clearly distinguish donor 82 contributions as both pre-treatment and control samples were assigned high human donor contribution percentages. Mice transplanted with human donor 82 had an average donor contribution percentage of 10.18% after transplant, while the pre-treatment samples had 11.95% and the control samples had 20.62% (Fig. 7c). SourceTracker also assigned very high donor contribution percentages to the pre-treatment and control samples in the study of double hu-mice transplanted with a mixture of all three human donors. Mice transplanted with the mixture of all three human donors had an average donor contribution percentage of 14.88% after transplant, while the pre-treatment samples had 21.04% and the control samples had 15.89% (Fig. 7d).

The high donor contribution percentages of donor 82 to the pre-treatment and control samples originated from an ASV with taxonomic assignment to *Akkermansia muciniphila.* This ASV was highly abundant in both hu-mice and double hu-mice and was much more prevalent in human donor 82 and the mixed donor sample compared to donors 65 or 74. To get a more accurate account of the stability of the human-like gut microbiome in post-transplant samples, we removed this ASV that was resulting in false positive donor contributions and once again used SourceTracker (Supplemental Figure - SourceTracker_82_Mix.pdf). After the removal of the ASV, the double hu-mice engrafted with human donor 82 had an average donor contribution percentage of 12.64% after transplant and double hu-mice engrafted with the mixture of all three human donors had an average donor contribution percentage of 18.11% after transplant, while all pre-treatment control samples were at 0.00%. Using both a percentage of shared ASVs and SourceTracker, we have demonstrated that double hu-mice had a stable human-like gut microbiome for the duration of the study, up to 14.5 weeks post­ transplant.

### Double hu-mice have increased human-like predicted metagenome functional content

In addition to evaluating microbiome classification, we also sought to assess the functional capacity of the microbiomes in double hu-mice. PICRUSt was used to predict the metagenome functional content from the 16S rRNA data after ASV inference [25], and the predicted KO features were graphed using both NMDS and PCoA plots (Fig. 8). Many of the double hu-mice samples clustered closer to the human donor samples than the hu-mice samples (Fig. 8a & 8b). When color-coded by donor and cohort, the microbiomes that clustered closest to the human donor samples belonged to mice from the second cohort of double hu-mice generated by engrafting bacteria from human donor 65 (Fig 8c & 8d). Similarly, several samples from the other double hu-mice cohorts also separated themselves from the hu-mice cluster and were closer to the human donor samples. (Fig. 8c & 8d).

**Figure 8.**
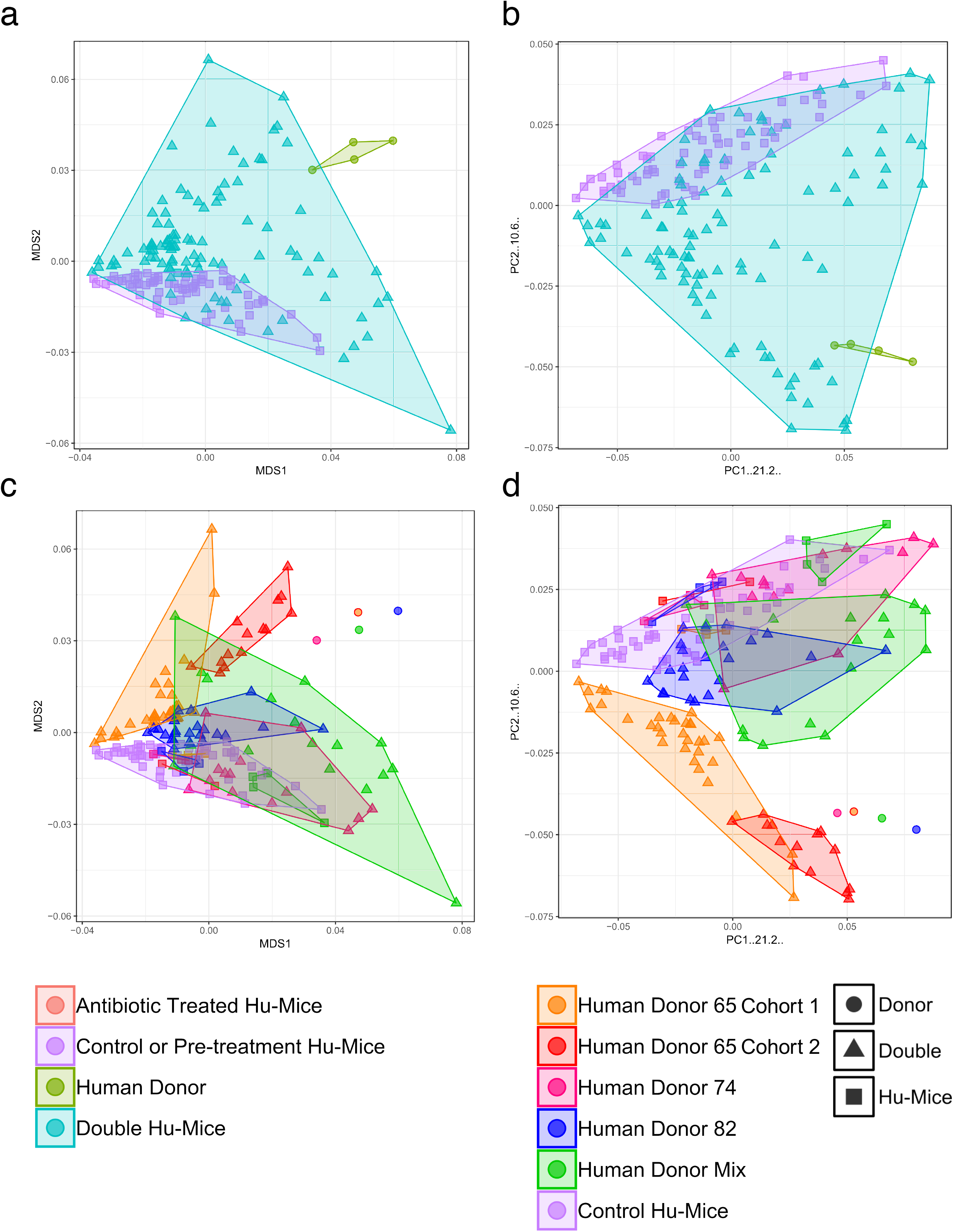
Double hu-mice have increased human-like predicted metagenome functional content. A) NMDS plot displaying the double hu-mice as a distinct cluster between the human fecal donor samples and pre-treatment or untreated control hu-mice. B) N MDS plot displaying human donor specific functional profiles in double hu-mice. C) PCoA plot displaying the double hu-mice as a distinct cluster between the human fecal donor samples and pre-treatment or untreated control hu-mice. D) PCoA plot displaying human donor specific functional profiles in double hu-mice.

We next tested for differences of each predicted KO feature among the three different groups of mice (Supplemental File - KO_Significant_Differences.xlsx). In total, there were 4,513 non-zero predicted KO features. Using Kruskal-Wallis testing and an FDR adjusted p-value of < 0.05, we found 35.54% (1,604/4,513) significantly different predicted KO features between hu­ mice and human donor samples. There were 39.09% (1,764/4,513) significantly different predicted KO features between double hu-mice and hu-mice samples. However, when we compared double hu-mice and the human donor samples, there were only 1.35% (61/4,513) significantly different predicted KO features. To clarify what functional aspects were missing from the double hu-mice gut microbiomes, we determined the predicted ASV contribution for each of the 61 significantly different KO features between the double hu-mice and human donor samples (Supplemental File - KO_Metagenome_Contribution.xlsx). This analysis provided insight into which functionally significant bacteria were not successfully engrafted from the human donor samples to the double hu-mice. A total of 95 ASVs with 73 unique taxonomic assignments were found to contribute to the 61 significantly different KO features (Supplemental Figure - KO_Contributions.pdf). Family level taxa with the highest levels of contribution included *Actinomycetaceae* (10.46%), *Bifidobacteriaceae* (8.76%), *Streptococcaceae* (20.74%), *Lachnospiraceae* (23.95%), and *Peptococcaceae* (8.66%). *Bifidobacterium adolescentis* was the highest contributing species (7.85%), and the highest contributing ASV (13.01%) had the taxonomic assignment of an unclassified *Streptococcus* species. Collectively, these analyses demonstrate that the predicted functional capacity of double hu-mice is more similar to human donor samples compared to hu-mice.

## Discussion

The goal of this study was to investigate the establishment and stability of an engrafted human gut microbiome after antibiotic pre-treatment in hu-mice. We call these mice double hu-mice as they have both a functional human immune system and human bacterial gut microbiomes similar to human donor samples. Our approach created highly reproducible and donor-specific human-like gut microbiomes across multiple cohorts of double hu-mice. Further, we showed that double hu-mice had increased measures of diversity and increased functional capacity compared to hu-mice. The engrafted human-like gut microbiomes were also stable for the duration of the study, up to 14.5 weeks post-transplant. Further, we demonstrated that the predicted functional capacity of double hu-mice is also more similar to the human donor samples than hu-mice.

One of the most significant aspects of the study was that the double hu-mice had gut microbiome profiles that were unique to the human donor engrafted. Each of the 4 different donor samples resulted in a distinct population of double hu-mice resembling the human donor. Double hu-mice create a huge opportunity for implementing personalized medicine and translational research. Potential applications for double hu-mice include testing for personalized responses of patient microbiomes to drug regimens, therapies, or dietary interventions. These mice could also be used to identify mechanisms underlying observations from human studies establishing a connection between the gut microbiome and disease.

In this study, we created a cohort of double hu-mice by engrafting a mixture of all three human donors to create an ‘un-biased’ human gut microbiome profile. Further study is needed to determine whether a mixed sample is beneficial in creating an un-biased human-like profile or if mixing samples creates an un-natural or unstable community of gut bacteria after engraftment. It may be advantageous to use a mixture of fecal samples derived from a large population to determine the broad impact of different treatments or diets to the human gut microbiome. This un-biased mix of a large population of donors could be used to add additional information to double hu-mice generated with individual donor profiles.

We found that double hu-mice had increased measures of alpha diversity compared to hu-mice. Several studies have highlighted the importance of microbiome diversity within the gut and have linked low gut microbiome diversity with several disease conditions[26, 27]. While not all low diversity conditions are detrimental, specifically when there is enrichment of potentially beneficial bacteria through prebiotic or probiotic treatment, the low pre-existing diversity found within the hu-mice was far below the levels observed in our human donor samples. After engraftment, the double hu-mice had increased levels of alpha diversity and maybe more importantly, had increased predicted functional capacities. This increased diversity may also offer a more realistic gut environment as it allows for diverse reciprocal interactions with the engrafted human immune system.

We also found that the engrafted human-like microbiome was very stable in our model for the length of the study, up to 14.5 weeks after transplant. We used several methods to determine the engraftment level and stability of the gut microbiome after transplant and found no reversion to the pre-existing murine profile. This stability allows study of the role of the gut microbiome in many human diseases such as HIV-1 and cancer. One outstanding question is whether the unique presence of human immune cells plays a role in stabilizing or enhancing engraftment of the human-like gut microbiome in our model compared to other non­ humanized mouse models. Our data showed no reversion to the pre-existing murine gut microbiome profile, perhaps due to some enhanced stability or selection by the reconstituted human immune system. Further studies are needed to determine the relationship between the engrafted gut microbiome and human immune system.

Many different methods and antibiotic regimens have been used for preconditioning of mice prior to fecal transplantation[28–30]. Different combinations and durations of antibiotic treatments may increase the efficiency of the fecal transplant into the host[29]. While the combination of Metronidazole, Ampicililn, Neomycin, and Vancomycin is widely used due to its broad spectrum of bacterial targets, the best methods are still being investigated. We found that the very rigorous method of gavaging antibiotics twice daily for 14 days used by Hintze et al. was too invasive for our NSG hu-mice and resulted in increased mortality[31]. Providing the antibiotics in the drinking water proved to be less stressful with improved health and survival of the mice. Meanwhile, we also found that 14 days was the optimum duration of antibiotic pre­ treatment to generate double hu-mice.

As expected, there is not a complete reconstitution of the human fecal donor profile in our double hu-mice due to several hypothesized reasons. There are major differences between the human and mouse digestive tract including structure, function, and pH [19]. Our mice are not germ-free to begin with, and the broad-spectrum antibiotic treatment can only reduce the prevalence of pre-existing murine gut bacteria. There were several key differences in the reconstituted mice compared to the human donors. Double hu-mice had significantly lower levels of several types of *Clostridia* including *Lachnospiraceae, Blautia, Coprococcus, Roseburia, Faecalibacterium,* and *Ruminococcus* compared to human donor samples. Many of these bacteria are well documented to be difficult to reconstitute within germ-free and SPF mouse models[20, 31, 32]. Similar to fecal transplants in humans designed to treat C. *difficile* infections, the engrafted human-like gut microbiome in our double hu-mice is the result of a combination of host, donor, and environmental bacteria[33]. Despite these previously known limitations, our double hu-mice model reproducibly results in a donor specific, stable, human­ like gut microbiome in the presence of a human immune system.

Germ-free animals are the gold standard for studying the gut microbiome. Using germ­ free mice to study the impact of the gut microbiome has been well-documented[l, 20). Germ­ free animal models may allow for a more complete reconstitution of a human-like gut microbiome following fecal transplant, however these models often do not have human immune system. Studying human immune reconstitution in hu-mice and pathogenesis of human specific diseases in a gnotobiotic environment could reveal important clues about the role of the gut microbiome. Nevertheless, many important mouse strains are not commercially available as germ-free, including NSG mice. Several studies have also shown that gnotobiotic mice may have long-lasting immune deficiencies, even after gut microbiome reconstitution[34–36]. Further, working with germ-free animals requires gnotobiotic facilities and equipment that is expensive and has limited availability. Lundberg et al., and Kennedy et al., nicely review the advantages and disadvantages of using antibiotic-treated versus germ-free rodents for microbiota transplantation studies [37, 38].

Our double hu-mice have the advantage of requiring only SPF housing conditions, which are widely available and less expensive compared to germ-free facilities. It also does not perturb the complex surgical procedures in generating BLT hu-mice because there is no need for a completely germ-free environment. Our NSG mice are housed and bred under SPF conditions and the diversity of their murine gut microbiota is low. Their immunodeficiency may contribute to their pre-existing low diversity gut microbiome status before human fecal material transplant. In a study by Zhou et al., NSG and C57BL6/J mice whose native microbiota were depleted by antibiotics followed by FMT had no significant differences in diversity but did observe significant differences in which species colonized[39]. A study done by Ericsson et al. showed that it is easier to transfer high diversity fecal donor samples into low diversity recipients, which could help to explain the success of engraftment and stability in our model[30]. Many questions remain as to the best antibiotic preconditioning regimen, the timing of fecal transplants, the total number of fecal transplants, the route of administration (oral versus rectal), the use of antacids, and preconditioning with osmotic laxatives such as polyethylene glycol, diet, and housing. Methods to optimize murine bacterial depletion along with reconstitution and stability of human specific bacteria are currently being explored.

## Conclusion

Here, we describe successful and stable transplantation of human fecal microbiomes into immunodeficient NSG mice surgically engrafted with a functional human immune system to create double hu-mice with human donor-specific human gut microbiomes. Double hu-mice will be beneficial to many applications of personalized medicine to test the impact of the human gut microbiome on human health and disease in the presence of a human immune system.

## Methods

### Generation of humanized BLT mice

All methods described here were conducted as we previously reported in accordance with Institutional Animal Care and Research Committee (IACUC)-approved protocols at the University of Nebraska-Lincoln (UNL)[22, 40–42]. The IACUC at the University of Nebraska­ Lincoln (UNL) has approved two protocols related to generating and using humanized BLT (hu­ BLT) mice, including Double Hu-Mice. Additionally, the Scientific Research Oversight Committee (SROC) at UNL has also approved the use of human embryonic stem cells and fetal tissues, which are procured from the Advanced Bioscience Resources for humanized mice studies (SROC# 2016-1-002).

Briefly, 6- to 8-week-old female NSG mice (NOD.*Cg-Prkdc^scid^IL2rg^tm1wjl^/SzJ*, catalog number 005557; (Jackson Laboratory) were housed and maintained in individual microisolator cages in a rack system capable of managing air exchange with prefilters and HEPA filters. Room temperature, humidity, and pressure were controlled, and air was also filtered. Mice were fed irradiated Teklad global 14% protein rodent chow (Teklad 2914) and were given autoclaved acidified drinking water. The second cohort of double hu-mice engrafted with fecal material from Donor 65 were supplemented with a high calorie gel (DietGel Boost). On the day of surgery, mice received whole-body irradiation at the dose of 12 cGy/gram of body weight with the RS200 X-ray irradiator (RAD Source Technologies, Inc., GA) and were then implanted with one piece of human fetal thymic tissue fragment sandwiched between two pieces of human fetal liver tissue fragments within the murine left renal capsule. Within 6 hours of surgery, mice were injected via the tail vein with 1.5 × 10^5^ to 5 × 10^5^ CD34^+^ hematopoietic stem cells isolated from human fetal liver tissues. Human fetal liver and thymus tissues were procured from Advanced Bioscience Resources (Alameda, CA). After 9 to 12 weeks, human immune cell reconstitution in peripheral blood was measured by a fluorescence-activated cell sorter (FACS) Aria II flow cytometer (BD Biosciences, San Jose, CA) using antibodies against mCD45-APC, hCD45-FITC, hCD3-PE, hCD19-PE/Cy5, hCD4-Alexa 700, and hCD8-APC-Cy7 (catalog numbers 103111, 304006, 300408, 302209, 300526, and 301016, respectively; Bio legend, San Diego, CA). Raw data were analyzed with FlowJo (version 10.0; FlowJo LLC, Ashland, OR). All mice used in this study had high levels of human immune cell reconstitution with an average of 85% hCD45+ cells in peripheral blood 10 weeks post-surgery. The mice were randomly assigned into experimental groups with similar immune reconstitution levels.

### Antibiotic treatment

A broad-spectrum antibiotic cocktail was prepared fresh daily consisting of Metronidazole (1 g/L), Neomycin (1 g/L), Vancomycin (0.5 g/L), and Ampicillin (1 g/L). The antibiotic cocktail was given to the mice ad libitum in the drinking water along with grape flavored Kool-Aid to improve palatability. Control group mice were given only grape flavored Kool-Aid in the drinking water. During antibiotic treatment, cages were changed daily to limit re-inoculation of pre-existing bacteria to the mice due to their coprophagic behavior. Antibiotics were given for 14 days for double hu-mice reconstituted with Donor 65 and Donor Mix and for 7 days in double hu-mice reconstituted with Donor 74 and Donor 82. Mice in the Pilot Study were given antibiotics via oral gavage. Mice were first given three days of anti-fungal Amphotericin B treatment (1 mg/kg) twice daily via oral gavage. Mice were then given the antibiotic cocktail along with Amphotericin B via twice daily via oral gavage. After 4 days of treatment, the Amphotericin B was stopped due to toxicity concerns and after 10 days of treatment oral gavage was reduced to once daily. Post-antibiotic treatment, mice were given autoclaved non-acidified deionized drinking water.

During the first few days of antibiotic treatment, the mice lost a considerable amount of body weight (10-20%). The weight loss plateaued at 3-4 days and remained steady for the remainder of antibiotic treatment. Body weight was carefully monitored during this time and If needed, mice were treated with Intraperitoneal (IP) injections of Ringer’s solution to mitigate any effects of dehydration. After fecal transplant, the mice began to regain weight and returned to their pre-existing weight within 2 weeks post-transplant. During antibiotic treatment, there was a large reduction in spleen size and a large increase in cecum size compared to controls. This is similar to the morphology observed in germ-free mice, providing further evidence for the efficacy of the antibiotic regimen[35].

### Donor samples and Fecal transplant

At 24 and 48 hours after the completion of antibiotic pre-treatment, mice were given 200 ul of human fecal material via oral gavage. OpenBiome supplied 3 ***FMT*** Upper Delivery Microbiota Preparations from 3 different healthy human donors (Donor 65, Donor 74, Donor 82). Samples were thawed once before fecal transplant to aliquot the samples within an anaerobic chamber. During this step, an equal portion of each of the samples were mixed together to create an unbiased human donor sample (Donor Mix). 16S rRNA sequencing data on the three donors was also supplied by OpenBiome (Supplementary data).

### Mouse fecal collection and DNA extraction

Individual mice were placed into autoclaved paper bags within a biosafety hood until fresh fecal samples were produced. Fecal samples were stored in 1.5 ml Eppendorf tubes at −80°C until DNA extraction. DNA was extracted from the fecal samples using the phenol:chloroform:isoamyl alcohol with bead beating method described previously [43]. Briefly, fecal samples were washed three times with 1 ml PBS buffer (pH 7). After the addition of 750 ul of lysis buffer, samples were transferred to tubes containing 300 mg of autoclaved 0.1 mm zirconia/silica beads (Biospec). 85 ul of 10% SDS solution and 40 ul of Proteinase K (15mg/ml, MC500B Promega) were added and samples were incubated for 30 minutes at 60^°^ C. 500 ul of Phenol:Chloroform:Isoamyl alcohol (25:24:1) was added and then samples were vortexed. Samples were then put into a bead beater (Mini-beadbeater 16 Biospec) for 2 minutes to physically lyse the cells. The upper phase of the sample was collected and an additional 500ul of Phenol:Chloroform:Isoamyl alcohol (25:24:1) was added. After samples were vortexed and spun down, the DNA in the upper phase was further purified twice with 500 ul of Phenol:Chloroform:Isoamyl alcohol (25:24:1). and was then precipitated with 100% Ethanol (2.5 × volume of sample) and 3M Sodium acetate (.1 × volume of sample) overnight at −20^°^ C. Samples are then centrifuged and dried at room temperature. DNA was resuspended in 100 ul of Tris-Buffer (10mM, pH8) and stored at −20^°^ C. DNA samples were quality checked by nanodrop (N D-1000 Nanodrop) .

### 16S rRNA gene sequencing

16S rRNA gene sequencing was performed at the University of Nebraska Medical Center Genomics Core Facility using xxx (detailed Illumina instrument here). DNA normalization and library prep were performed followed by V3-V4 16S rRNA amplicon gene sequencing using a MiSeqV2 (Illumina) The following primer sequences were used: (Primer sequences: Forward Primer = 5’

TCGTCGGCAGCGTCAGATGTGTATAAGAGACAGCCTACGGGNGGCWGCAG 16S Amplicon PCR

Reverse Primer = 5’

GTCTCGTGGGCTCGGAGATGTGTATAAGAGACAGGACTACHVGGGTATCTAATCC

Illumina overhangs: Forward overhang: 5’

TCGTCGGCAGCGTCAGATGTGTATAAGAGACAG-[locusspecific sequence] Reverse overhang: 5’ GTCTCGTGGGCTCGGAGATGTGTATAAGAGACAG-[locusspecific sequence]).

### Generation of the amplicon sequence variant table and data analysis

Illumina-sequenced paired-end fastq files were demultiplexed by sample and barcodes were removed by the sequencing facility. The University of Nebraska Holland Computer Center Crane cluster was used to run the DADA2 v1.8 R package in order to generate an amplicon sequence variant (ASV) table[44]. An example of the script used to generate the ASV table is provided in the supplementary materials. The DADA2 pipeline was performed as follows, sequences were filtered and trimmed during which any remaining primers, adapters, or linkers were also removed. The sequencing error rates were estimated using a random subset of the data. Dereplication of the data combined all identical sequencing reads into unique sequences with a corresponding abundance. The core sample inference algorithm was then applied to the dereplicated data. The forward and reverse reads were then joined to create the full denoised sequences and an initial ASV table was generated. Any sequences outside the expected length for the V3-V4 amplicon were then filtered from the table. Chimeric sequences were then removed and a final ASV table was generated. Taxonomy was assigned using the Greengenes 13.8 database and RDP Classifier with a minimal confidence score of 0.80[45, 46].

### Data analysis

Analysis was performed using R package mctoolsr (https://github.com/leffj/mctoolsr/). and samples were rarified to 13,000 ASVs for downstream analysis. Additional testing of differences between groups was performed using LEfSe[23]. SourceTracker was used to evaluate the stability of the transferred donor microbiome in the double hu-mice[24]. GraphPad Prism 5 were used to create some figures. DADA2 generated ASVs were used to predict the functional metagenome capacity using PICRUSt[25] via the following pipeline (https://github.com/vmaffei/dada2_to_picrust).

### Data availability

The datasets generated during the current study are available in the NCBI SRA repository, [https://www.ncbi.nlm.nih.gov/bioproject/PRJNA507247].

## Acknowledgements

We would like to thank Yanmin Wan, Guobin Kang, and Pallabi Kundu for their assistance in generating BLT hu-mice. We would like to acknowledge the UNMC Genomics Core Facility who receives partial support from the Nebraska Research Network In Functional Genomics N E-INBRE P20GM103427-14, The Molecular Biology of Neurosensory Systems CoBRE P30GM110768, The Fred & Pamela Buffett Cancer Center - P30CA036727, The Center for Root and Rhizobiome Innovation (CRRI) 36-5150-2085-20, and the Nebraska Research Initiative. We would like to thank University of Nebraska-Lincoln Life Sciences Annex and their staff for their assistance. This study is supported in part by the National Institutes of Health (NIH) Grants R01AI124804 (to Javis), R33AI122377 (Planelles), P30 MH062261-16A1 Chronic H IV Infection and Aging in NeuroAIDS (CHAIN) Center (to Buch & Fox), 1R01AI111862 and R21 AI143405 to Q Li. The funders had no role in study design, data collection and analysis, preparation of the manuscript or decision for publication.

## Author contributions

LD and QL designed the experiments and wrote the manuscript. LD performed experiments and analyzed the data. ART provided input on experimental design and manuscript preparation.

