## Supplementary figures and images for "Stable Engraftment of a Human Gut Bacterial Microbiome in Double Humanized BLT-mice"

### Supplemental Data 1

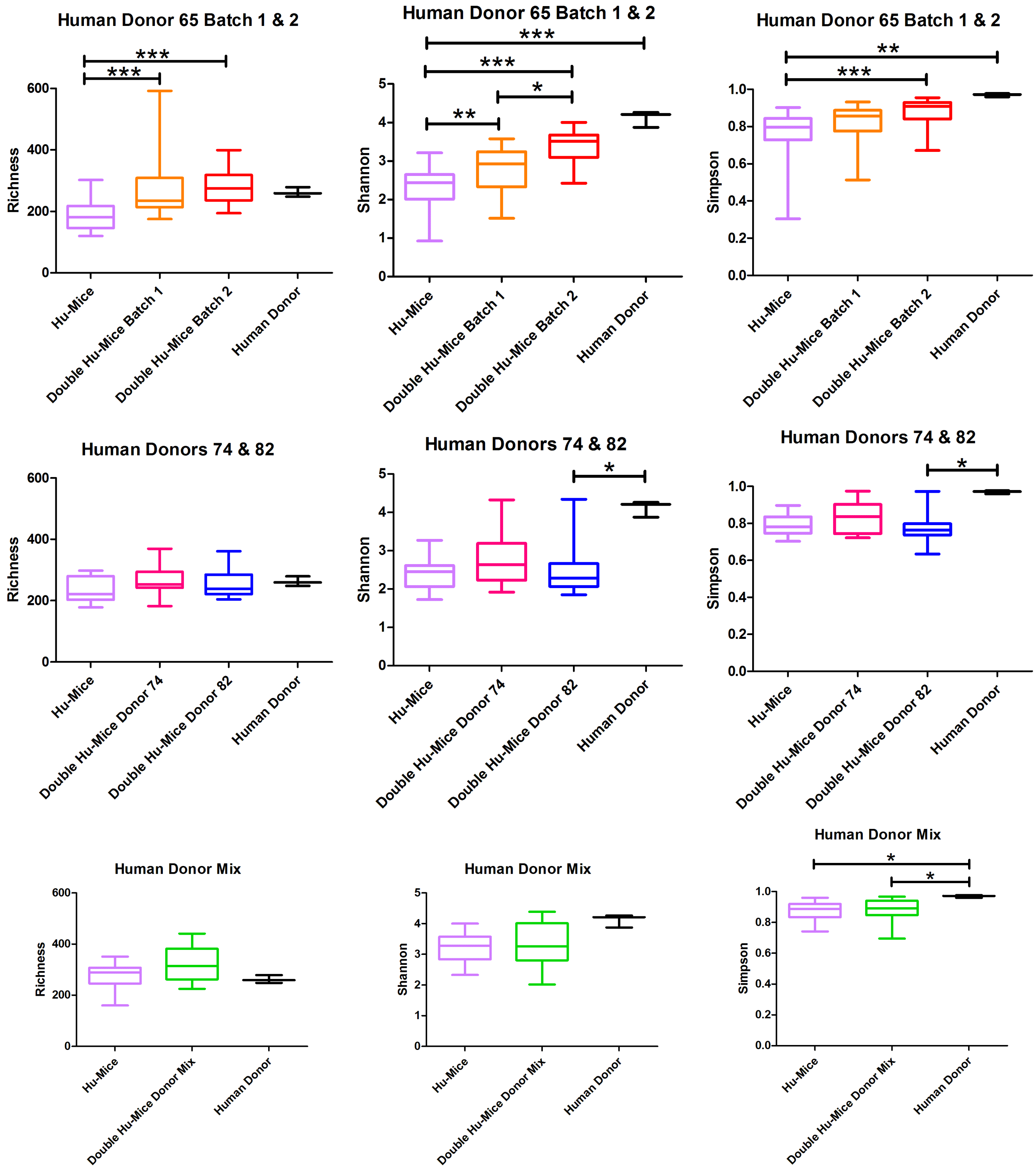

### Supplemental Data 2

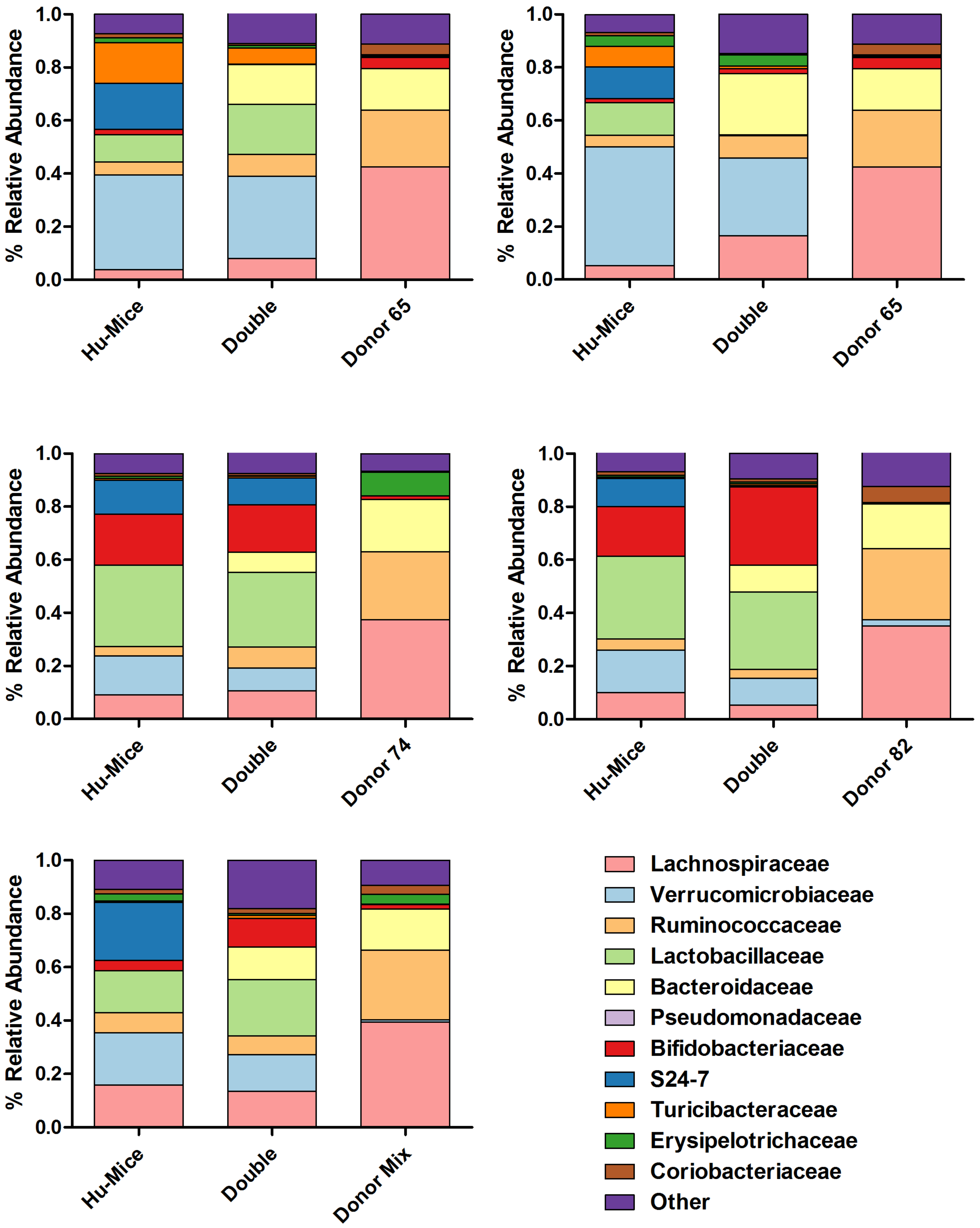

### Supplemental Data 5

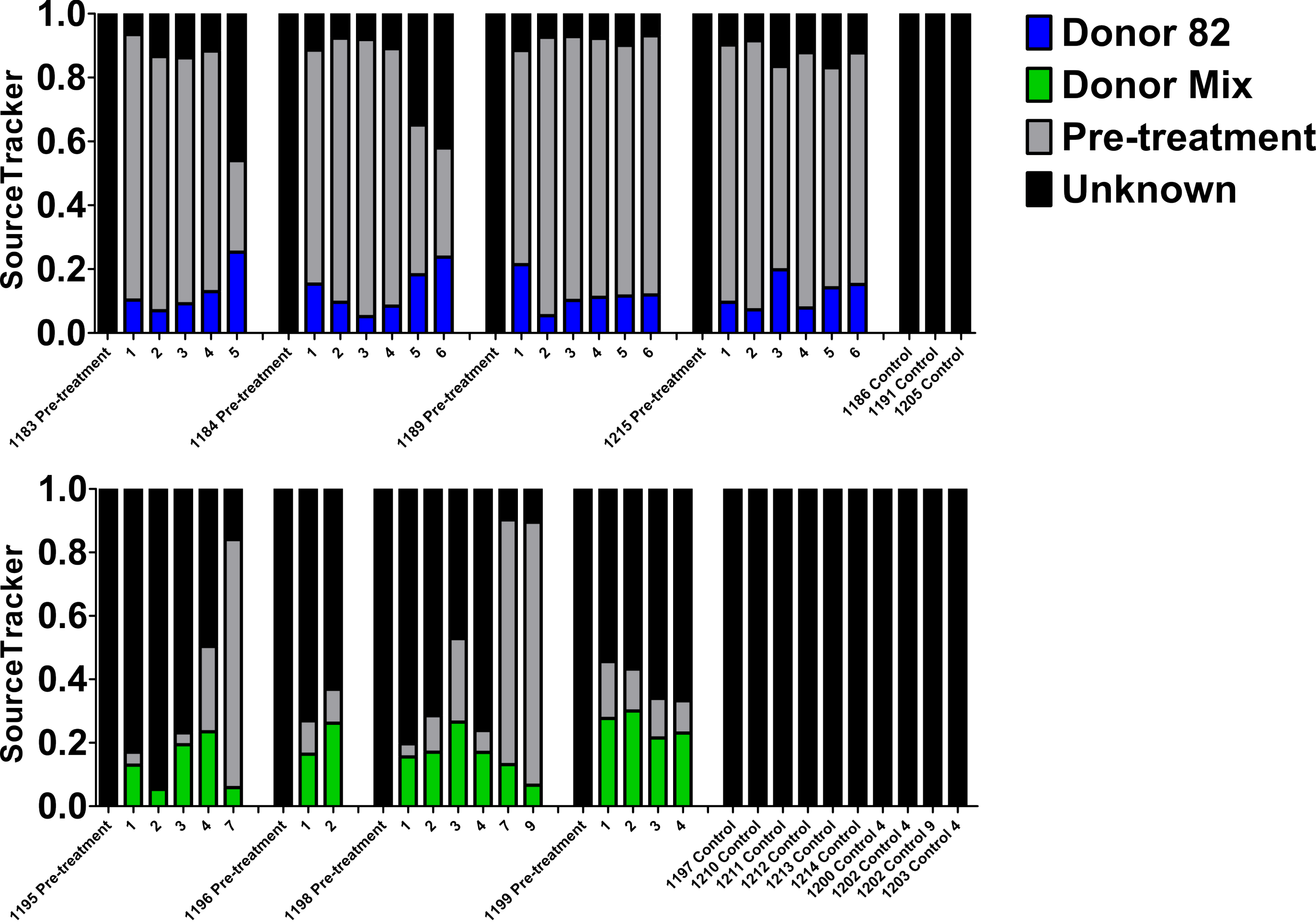

### Supplemental Data 8

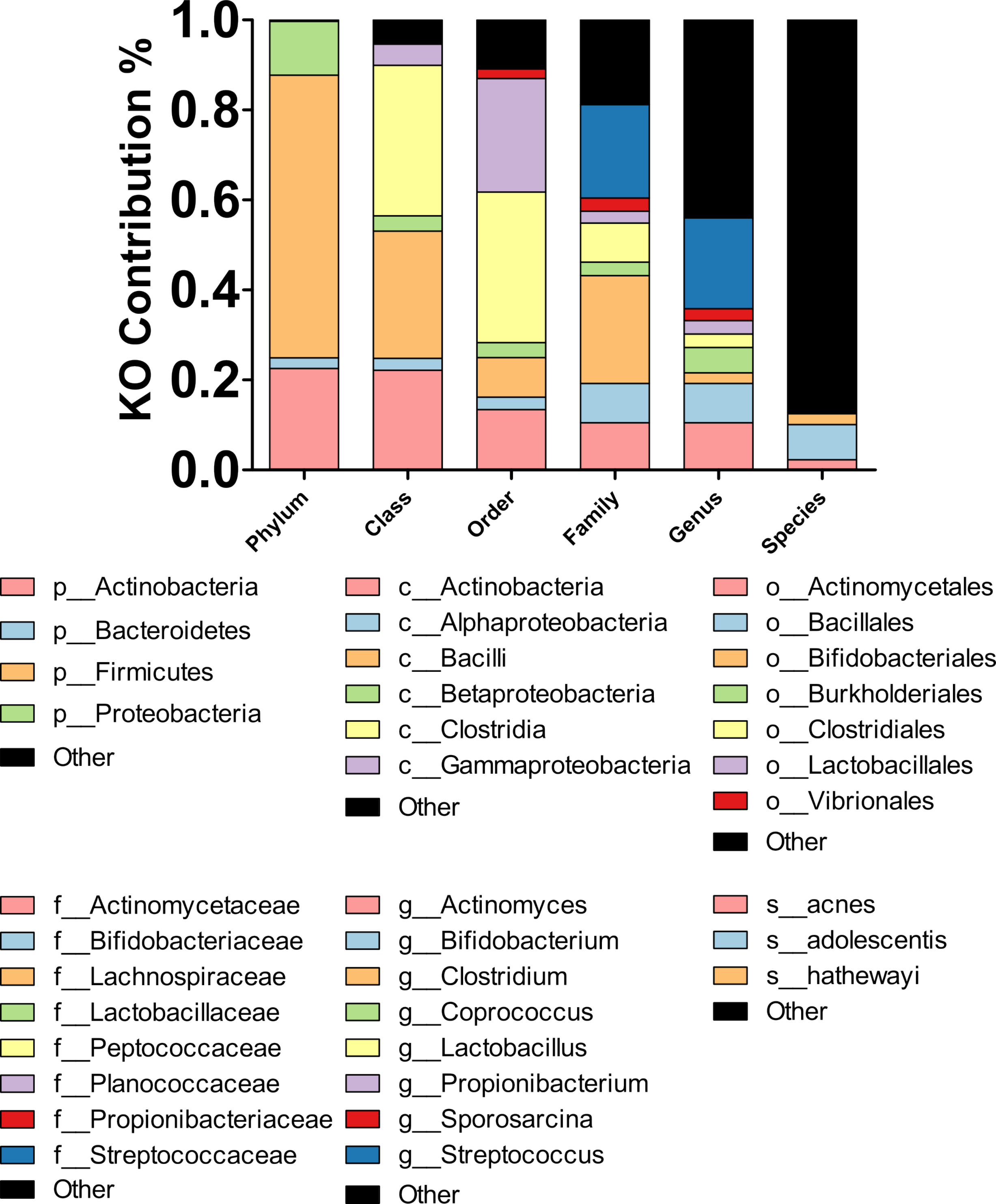
